# Digesting the data: Proper validation in ancient metagenomic studies is essential

**DOI:** 10.1101/2024.02.27.581519

**Authors:** Aleksandra Laura Pach, Liam T Lanigan, Jonas Niemann, Mikkel Winther Pedersen, Hannes Schroeder

**Affiliations:** Globe Institute, Faculty of Health and Medical Sciences, University of Copenhagen, Øster Farimagsgade 5A, 1253 Copenhagen, Denmark; School of Archaeology, University of Copenhagen, Karen Blixens Plads 8, 2300 Copenhagen Denmark

**Author notes:** correspondence (H.S.).

## Abstract

In a recent publication in this journal, Reynoso-García et al. [1] used shotgun sequencing to analyze human coprolites (paleofeces) from two pre-Columbian contexts in Puerto Rico to reconstruct the diet of the island’s Indigenous population before the arrival of Europeans. Based on the results, the team claim to have identified various edible New World plant species, including maize (*Zea mays*), sweet potato (*Ipomoea batatas*), chili pepper (*Capsicum annuum*), peanut (*Arachis spp*.), papaya (*Carica papaya*), and tomato (*Solanum lycopersicum*), as well as other cultivars such as cotton (*Gossypium barbadense*) and tobacco (*Nicotiana sylvestris*) [1]. Reynoso-García et al. [1] also claim to have identified edible fungi, including *Ustilago spp*., which according to the authors, further supports their findings and points to the consumption of *huitlacoche* or corn smut, a known delicacy in Mexico today that is believed to have originated in Aztec times [2].

Shotgun DNA sequencing of archaeological samples, such as dental calculus or coprolites, provides a powerful tool to reconstruct ancient microbial communities and to study the evolution of the human microbiome [e.g. 3,4–6]. In some instances, shotgun sequencing results can also provide insights into the diet and subsistence strategies of past communities [e.g. 7,8]. However, identifying DNA from dietary sources in complex shotgun metagenomic datasets is far from straightforward. As has been discussed previously [9,10], and as we demonstrate below, one of the main challenges is the risk of false positives. Any potential dietary signals should, therefore, be carefully assessed. Unfortunately, we feel that Reynoso-García et al.’s study [1] falls short in that regard and we conclude that while it is entirely possible (and even likely, based on other evidence [e.g. 11,12]) that the Indigenous inhabitants of Puerto Rico subsisted on a diet that included some, or even all of the edible plant taxa the team identified, the DNA results they present do not, in and of themselves, support that claim.

## Lack of validation casts doubt on results

Paleodietary reconstruction using ancient metagenomics is challenging for several reasons [9,10]. As Reynoso-García et al. [1] rightly point out, one of the main challenges is the risk of modern contamination and it is essential to prevent and control for DNA contamination from the field to the laboratory [13,14]. However, controlling for present-day contamination during sampling and in the laboratory alone is not enough as false positive results can arise for other reasons, including sequence homology or the presence of contamination in published reference genomes [9,15]. Therefore, any potential dietary hits should be independently verified using additional analyses. Some of the features that should be assessed include the similarity of the reads to the reference, randomness of genomic coverage, and the presence of ancient DNA damage patterns. All these criteria have been widely employed in the field of ancient metagenomics to confirm true positive hits, e.g. to ancient pathogens [16–21]. Unfortunately, Reynoso-García et al. [1] did not employ any of these standards, making it difficult to judge the validity of the results.

One of the main problems with Reynoso-García et al.’s study [1] is their choice of taxonomic classifier. Of the many available tools, the team opted for Kajiu v1.5.0 [22], which, similar to DIAMOND [23], classifies DNA reads by first translating them into amino acid (AA) sequences. These are then compared against a protein reference database (e.g. NCBI RefSeq or NCBI’s non-redundant (nr) database) to find maximum exact matches. This approach makes Kaiju extremely memory-efficient and, therefore, typically faster than common k-mer-based methods such as Kraken [24] and Clark [25]. However, as Mann et al. [9] have pointed out, using AA sequences has several drawbacks. First, it limits database searches to the protein-coding regions of the genome only, reducing the number of potential hits. Second, typically only high-quality genomes are included in protein databases, reducing the number of possible taxa that can be identified. Third, short ancient DNA molecules translate into even shorter AA sequences that may match many different proteins, potentially resulting in less specificity [26] and higher rates of false positive hits when compared to DNA. In addition, others have shown that Kaiju’s sensitivity is low, particularly when the DNA fragments are short and damaged [27]. Another drawback is that by using AA sequences Kaiju [22] ignores important sequence information, such as ancient DNA damage patterns that are crucial for validating ancient DNA results [28]. Therefore, we feel that while tools like Kaiju [22] and DIAMOND [23] have been used for the classification of ancient metagenomic shotgun reads in the past [e.g. 5,29], there are now better tools available.

Following the initial classification with Kajiu (v1.5.0) [22], Reynoso-García et al. [1] used Meta-SourceTracker [30] to “verify the authenticity of coprolite DNA” and quantify the level of contamination in their samples. Meta-SourceTracker [30] uses Bayesian methods to estimate the relative contribution of samples with a known composition (sources) to a set of samples of unknown composition (sinks). In their study, Reynoso-García et al. [1] used a set of publicly available fecal, skin, and soil metagenomes as sources and the two pre-Columbian coprolites as sinks. The data were processed using Kaiju [22] and then used as input for Meta-SourceTracker [30] to estimate the relative proportion of each potential source to the archaeological samples. However, the team only used the taxonomic abundances for the eukaryotic domain as input, thereby ignoring most of the data (i.e. the microbial portion). By excluding the microbial hits, Reynoso-García et al. [1] ignore the portion of the data that is arguably most informative regarding the origin of a sample and that Meta-SourceTracker [30] uses to estimate source proportions of complex metagenomic sink samples and to assess the level of contamination. We therefore consider their use of Meta-SourceTracker [30] to be somewhat unconventional.

To further support their conclusions, Reynoso-García et al. [1] analyzed reads from phytopathogenic fungi, suggesting that the fungi could have been ingested alongside plants as part of the diet. Using network analysis, the team examined the relationships between specific fungal pathogens (e.g. *Ustilago* spp.) and plants they detected in the coprolites, suggesting that they may have been consumed together. What’s more, they use the presence of *Ustilago* spp. to suggest that the Indigenous inhabitants of Puerto Rico may have consumed *huitlacoche* or corn smut, a common fungal pathogen that is still consumed as a delicacy in Mexico and other parts of Mesoamerica today. Modeling phytopathogenic fungi and plant interactions is a very interesting idea that can potentially inform our understanding of past diets and cultural practices. However, many of the fungal species the team identified occur naturally in the environment. For example, *Ustilago* spp. are associated with grasses in general [31] and, unfortunately, it is impossible to determine–based on the data they present–whether the fungal reads Reynoso-García et al. [1] identified are ancient or whether they are present-day contaminants. Our reanalysis of the data suggests that they do not show any of the telltale signs of ancient DNA (short average fragment lengths and ancient DNA damage), suggesting that they are likely modern environmental contamination (Fig 1).

**Fig 1.**
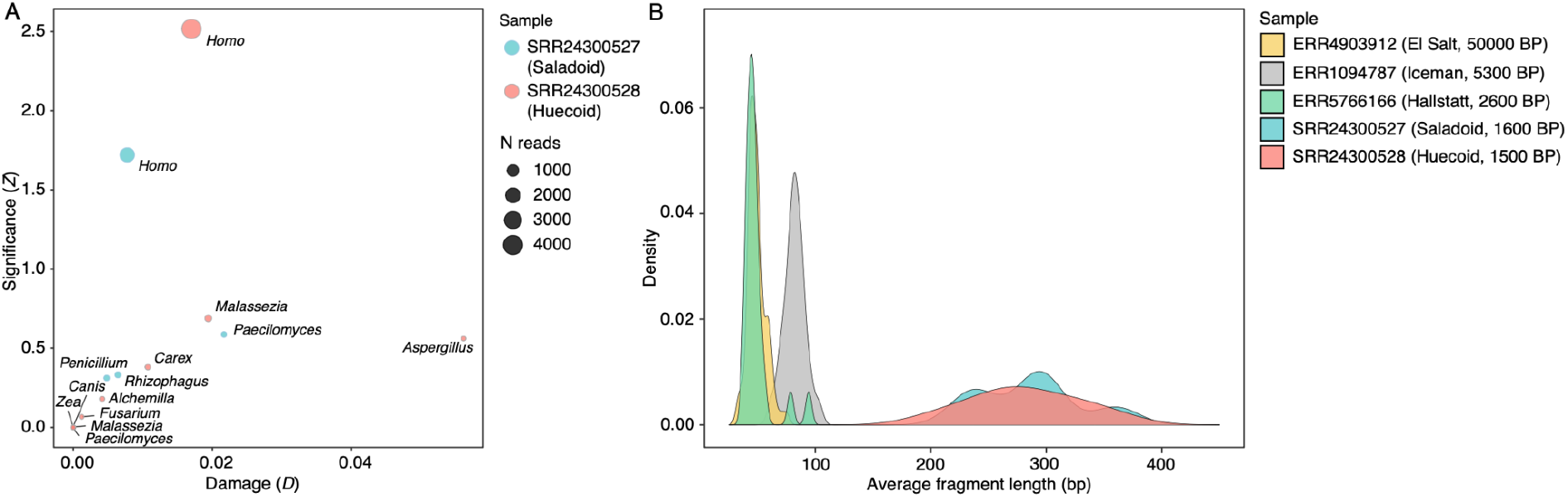
DNA damage and DNA fragment length distribution. **(A)** MetaDMG [32] statistics for eukaryotic taxa with >10 assigned reads detected in the two pre-Columbian coprolites sequenced by Reynoso-García et al. [1]. The Damage (*D*) statistic is an estimate of a background-substracted C-T damage frequency at the first read position, while the Significance (*Z*) statistic estimates how well the data fit the damage model (i.e. more damage at the beginning of the read) compared to the null model (i.e. constant amount of damage) with higher *Z* values indicating a better fit. **(B)** Density plot showing the average fragment length distribution of all eukaryotic reads detected (for taxa with >10 assigned reads) in the two pre-Columbian samples [1] and three samples from prehistoric contexts in Europe [8,33,34], for comparison. Dates are approximate sample ages in calibrated years BP (before present).

To try to validate their results, we parsed their data through a pipeline that was specifically designed for the classification and validation of ancient eukaryotic shotgun reads [35]. We first dereplicated the raw sequencing reads using vsearch v2.22.1 [36] running --fastx_uniques, --minseqlength 30, and --strand both. We then mapped all the reads against the RefSeq (release 213), NCBI nt (downloaded September 2022), and PhyloNorway database (September 2022) and parsed all merged and sorted alignments with at least 10 classified reads and 95% similarity to the reference to metaDMG v.1.17 [32] for taxonomic profiling and DNA damage estimation. All code, data and result files are available on GitHub (https://github.com/AleksandraLaura/DietComment). Using this workflow, we obtained 66,293 (SRR24300527) and 170,746 (SRR24300528) hits to dozens of eukaryotic taxa for the two Caribbean samples (S1 Table). However, none of the classified reads show any significant amount of ancient DNA damage (Fig 1A) suggesting that the DNA is not ancient. This is further supported by the fact that the average fragment length of all eukaryotic reads is 270 bp (Fig 1B), which far exceeds the fragment length typically observed in ancient samples, especially in the tropics [37–42].

Methodological shortcomings aside, the main problem with Reynoso-García et al.’s study [1] is that it is based on very little data. While it was difficult to assess how many reads their analysis is based on (as the authors report proportions rather than the actual number of reads), our reanalysis of the data shows that their results are based on very few reads, even single reads in some cases (S2 Table). This is problematic because validating true positive hits requires a minimum amount of data [9,43]. For example, several studies have shown that quantifying ancient DNA damage patterns requires at least 200-300 reads [9,32,43]. We recovered substantially more eukaryotic reads using our workflow than Reynoso-García et al. did with Kaiju [22], including several hundred hits to different plant and fungal taxa (S1 Table). This is expected as we are not only recovering matches to protein-coding regions of the genome but hits to the entire genome. However, as shown in Fig 1A none of these taxa show any significant amount of ancient DNA damage (*Z*>2; *D*>0.05), suggesting that they are likely not ancient [32]. So, where do these reads come from?

## How can false positives arise?

False positives are a common problem in ancient metagenomics and they can occur for several reasons including modern contamination from the burial environment (which will lack the characteristic ancient DNA damage patterns) or false alignments (which may well look ancient but are false positives nonetheless) [9,43]. The latter can occur for several reasons, including the presence of contamination in published reference genomes, sequence homology between related taxa, or the uneven taxonomic composition of reference databases [15,44]. For example, Steingegger and Salzberg identified over 2 million contaminated entries in GenBank affecting over 6,000 taxa [15]. This can easily lead to false positive alignments and eukaryotic genomes seem to be particularly affected due to their larger genome size and repetitive content (as compared to prokaryotes) [15]. While several methods for removing contamination have been developed, they are only now being implemented [15,45,46]. However, even with cleaned, well-annotated genomes, conserved regions and regions of low complexity can result in false positive alignments. This is especially true for higher eukaryotic taxa, with their larger genomes and many conserved regions across the genome [9].

To highlight the risk of false positives, we replicated Reynoso-García et al.’s workflow and applied it to three previously published datasets from prehistoric Europe [8,33,34]. Like Reynoso-García et al. [1], we first processed the raw data with metaWRAP (v. 1.3.2) [47] to remove adapters, poor-quality sequences, as well as human host DNA sequences. We then assembled a custom Kaiju (v. 1.9.2) [22] database using all the taxIDs from the provided Kaiju “nr_euk” database in addition to all Viridiplantae (taxID 33090). Lastly, we processed the reads with Kaiju as described in Reynoso-García et al. [1]. Results were plotted in R (v. 4.3.1) using ggplot2 (v. 3.4.3) and webr (v. 0.1.6). All source code and necessary files are available on GitHub (https://github.com/AleksandraLaura/DietComment). Worryingly, we detect the same nine New World species in the prehistoric European samples that Reynoso-García et al. [1] detected in the Caribbean coprolites, including maize (*Zea mays*), chili pepper (*Capsicum annuum*), papaya (*Carica papaya*), sweet potato (*Ipomoea batatas*), peanuts (*Arachis spp*.), and tomato (*Solanum lycopersicum*), as well as cotton (*Gossypium barbadense*) and tobacco (*Nicotiana sylvestris*) (Fig 2A)–a result that is difficult to explain unless we think that those taxa made it to Europe in pre-Columbian times.

**Fig 2.**
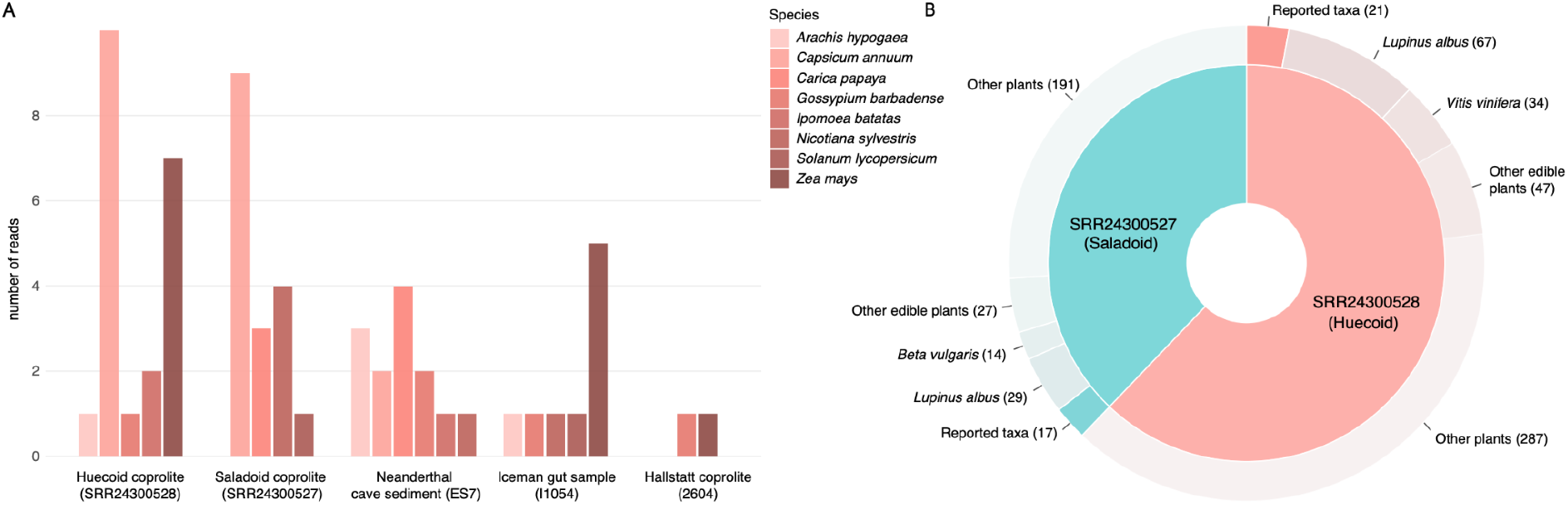
**(A)** Barplot showing the number of reads assigned to New World plant taxa in the two pre-Columbian coprolites from Puerto Rico [1], a c. 50,000-year-old sediment sample (ES7) from a Middle-Palaeolithic site in Spain [34], a gastrointestinal sample (I1054) from “Ötzi”, the Iceman who lived around 5,300 years BP [33], and a coprolite sample (2604) from the Hallstatt salt mine in Austria dating to around 2,600 years BP using Reynoso-García et al.’s [1] workflow. **(B)** Donut plot showing the number of reported and unreported plant taxa detected in the two Caribbean coprolite samples (SRR24300527 and SRR24300528) analyzed by Reynoso-García et al. [1]. The number of reads per taxon are given in parentheses.

We also reanalyzed Reynoso-García et al.’s data and found the same taxa they report (Fig 2B and <u>S2 Table</u>). However, in addition, we obtained hits to several taxa the team did not report, including grape (*Vitis vinifera*), white lupin (*Lupinus albus*), and beet (*Beta vulgaris*), which are all “Old World” taxa (Fig 2B and <u>S2 Table</u>) and it seems unlikely that they, too, formed part of the Puerto Rican diet in precolonial times. To try and determine where the plant reads Reynoso-García et al. [1] reported may have come from, we tried to collapse the reads and blasted [48] the resulting sequences against NCBI’s nt database. We find that most of the reads returned bacteria as the top hit or did not match anything in the database (<u>S3 Table</u>). Only nine reads returned maize (*Z. mays*) or tomato (*S. lycopersicum*) as the top hit. However, most of those hits are either partial (<50% query cover) or poor-quality (<90% identity) matches, or they are ambiguous, i.e. one of the reads returned a different taxon (e.g. *Sorghum bicolor*) as the top hit (<u>S3 Table</u>). Only three reads returned maize or tomato as an unambiguous hit. However, all of these fragments are uncharacteristically long for ancient DNA (>350 bp) and none of them show any signs of ancient DNA damage (in the form of C to T transitions), suggesting that they are not ancient.

## Validation remains essential for ancient metagenomic studies

While it is entirely possible and even likely based on other evidence [e.g. 11,12] that people in the precolonial Caribbean consumed maize, cassava, sweet potato and some of the other plant taxa that Reynoso-García et al. [1] identified in the two coprolites, the DNA evidence they present does not in and of itself support that claim. Based on our reanalysis of the data, it seems more likely that the hits they obtained are false positives resulting from false alignments [9]. This is supported by the length of the DNA fragments, which far exceeds the average fragment length of ancient DNA of similar age from the region [39–42]. A BLAST analysis of the Kaiju-classified reads suggests that the majority of them are bacterial. Alternatively, some of them could have derived from related taxa (e.g. grasses) that match conserved regions in the genome, which may explain some of the ambiguous BLAST hits. We have also shown that you can obtain hits to the same taxa (i.e. maize, cassava, sweet potato, etc.) when applying Reynoso-García et al.’s [1] workflow to a set of samples from prehistoric contexts in Europe, highlighting the risk of false positive results when using a metagenomics approach [9,10].

Given the risk of false positives, it is essential to assess the validity of the results of ancient metagenomic studies [9,17,20,43,49–51]. Generally, this involves at least two steps. First, it has to be demonstrated that the reads have been assigned to the correct taxon. This can be achieved by checking aspects of the data, such as the edit distance (ED) distribution or average nucleotide identity (ANI) scores of the alignments, which provide a measure of nucleotide-level genomic similarity between the reads and the reference. In addition, it should be verified that the reads are randomly distributed across the reference genome to check for spurious alignments to contaminated or conserved regions within the reference. Read pileups in specific regions of the genome are almost certainly false alignments that should be ignored or investigated further. Second, it should be demonstrated that the reads are indeed ancient by assessing the length of the recovered DNA fragments and the presence of ancient DNA damage patterns (in the form of C to T miscoding lesions) [28,32,52–56]. Without these checks, it is impossible to assess the results of any ancient DNA study, as likely or spectacular as they may be [9,57].

## Acknowledgments

We thank Tom Gilbert and Fernando Racimo for commenting on an earlier version of this manuscript.

## Supporting Information

[S1 Table: metaDMG results for the eukaryotic hits with at least 10 reads](https://github.com/AleksandraLaura/DietComment/blob/main/S1_Table.xlsx).

[S2 Table. Kaiju v. 1.9.2 results for two pre-Columbian coprolites and three previously published samples from prehistoric Europe](https://github.com/AleksandraLaura/DietComment/blob/main/S2_Table.xlsx).

[S3 Table: Top BLAST results for the New World plant reads identified by Kaiju in the Caribbean samples](https://github.com/AleksandraLaura/DietComment/blob/main/S3_Table.xlsx).

## References

1. Reynoso-García J, Santiago-Rodriguez TM, Narganes-Storde Y, Cano RJ, Toranzos GA. Edible flora in pre-Columbian Caribbean coprolites: Expected and unexpected data. PLoS One. 2023;18: e0292077.

2. Valverde ME, Hernández-Pérez T, Paredes-Lopez O. Huitlacoche – A 21^st^century culinary delight originated in the Aztec times. ACS Symposium Series. Washington, DC: American Chemical Society; 2012. pp. 83–100.

3. Fellows Yates JA, Velsko IM, Aron F, Posth C, Hofman CA, Austin RM, et al. The evolution and changing ecology of the African hominid oral microbiome. Proc Natl Acad Sci U S A. 2021;118. doi:10.1073/pnas.2021655118

4. Granehäll L, Huang KD, Tett A, Manghi P, Paladin A, O’Sullivan N, et al. Metagenomic analysis of ancient dental calculus reveals unexplored diversity of oral archaeal Methanobrevibacter. Microbiome. 2021;9: 197.

5. Ottoni C, Borić D, Cheronet O, Sparacello V, Dori I, Coppa A, et al. Tracking the transition to agriculture in Southern Europe through ancient DNA analysis of dental calculus. Proc Natl Acad Sci U S A. 2021;118. doi:10.1073/pnas.2102116118

6. Wibowo MC, Yang Z, Borry M, Hübner A, Huang KD, Tierney BT, et al. Reconstruction of ancient microbial genomes from the human gut. Nature. 2021. doi:10.1038/s41586-021-03532-0

7. Jensen TZT, Niemann J, Iversen KH, Fotakis AK, Gopalakrishnan S, Vågene ÅJ, et al. A 5700 year-old human genome and oral microbiome from chewed birch pitch. Nat Commun. 2019;10: 5520.

8. Maixner F, Sarhan MS, Huang KD, Tett A, Schoenafinger A, Zingale S, et al. Hallstatt miners consumed blue cheese and beer during the Iron Age and retained a non-Westernized gut microbiome until the Baroque period. Curr Biol. 2021;31: 5149–5162.e6.

9. Mann AE, Fellows Yates JA, Fagernäs Z, Austin RM, Nelson EA, Hofman CA. Do I have something in my teeth? The trouble with genetic analyses of diet from archaeological dental calculus. Quat Int. 2020. doi:10.1016/j.quaint.2020.11.019

10. Shillito L-M, Blong JC, Green EJ, van Asperen EN. The what, how and why of archaeological coprolite analysis. Earth Sci Rev. 2020;207: 103196.

11. Mickleburgh HL, Pagán-Jiménez JR. New insights into the consumption of maize and other food plants in the pre-Columbian Caribbean from starch grains trapped in human dental calculus. J Archaeol Sci. 2012;39: 2468–2478.

12. Siegel PE, Pearsall DM. Plant resource diversity in the ethnobotanical record of precolonial Puerto Rico: Evidence from microbotanical remains. J Archaeol Sci. 2023;160: 105859.

13. Llamas B, Valverde G, Fehren-Schmitz L, Weyrich LS, Cooper A, Haak W. From the field to the laboratory: Controlling DNA contamination in human ancient DNA research in the high-throughput sequencing era. STAR: Science & Technology of Archaeological Research. 2017;3: 1–14.

14. Peyrégne S, Prüfer K. Present-Day DNA Contamination in Ancient DNA Datasets. Bioessays. 2020;42: e2000081.

15. Steinegger M, Salzberg SL. Terminating contamination: large-scale search identifies more than 2,000,000 contaminated entries in GenBank. Genome Biol. 2020;21: 115.

16. Key FM, Posth C, Krause J, Herbig A, Bos KI. Mining Metagenomic Data Sets for Ancient DNA: Recommended Protocols for Authentication. Trends Genet. 2017;33: 508–520.

17. Hübler R, Key FM, Warinner C, Bos KI, Krause J, Herbig A. HOPS: automated detection and authentication of pathogen DNA in archaeological remains. Genome Biol. 2019;20: 280.

18. Spyrou MA, Bos KI, Herbig A, Krause J. Ancient pathogen genomics as an emerging tool for infectious disease research. Nat Rev Genet. 2019;20: 323–340.

19. Duchêne S, Ho SYW, Carmichael AG, Holmes EC, Poinar H. The Recovery, Interpretation and Use of Ancient Pathogen Genomes. Curr Biol. 2020;30: R1215–R1231.

20. Der Sarkissian C, Velsko IM, Fotakis AK, Vågene ÅJ, Hübner A, Fellows Yates JA. Ancient Metagenomic Studies: Considerations for the Wider Scientific Community. mSystems. 2021;6: e0131521.

21. Sikora M, Canteri E, Fernandez-Guerra A, Oskolkov N, Ågren R, Hansson L, et al. The landscape of ancient human pathogens in Eurasia from the Stone Age to historical times. bioRxiv. 2023. p. 2023.10.06.561165. doi:10.1101/2023.10.06.561165

22. Menzel P, Ng KL, Krogh A. Fast and sensitive taxonomic classification for metagenomics with Kaiju. Nat Commun. 2016;7: 11257.

23. Buchfink B, Xie C, Huson DH. Fast and sensitive protein alignment using DIAMOND. Nat Methods. 2015;12: 59–60.

24. Wood DE, Salzberg SL. Kraken: ultrafast metagenomic sequence classification using exact alignments. Genome Biol. 2014;15: 1–12.

25. Ounit R, Wanamaker S, Close TJ, Lonardi S. CLARK: fast and accurate classification of metagenomic and genomic sequences using discriminative k-mers. BMC Genomics. 2015;16: 236.

26. Eisenhofer R, Weyrich LS. Assessing alignment-based taxonomic classification of ancient microbial DNA. PeerJ. 2019;7: e6594.

27. Pusadkar V, Azad RK. Benchmarking Metagenomic Classifiers on Simulated Ancient and Modern Metagenomic Data. Microorganisms. 2023;11. doi:10.3390/microorganisms11102478

28. Dabney J, Meyer M, Pääbo S. Ancient DNA damage. Cold Spring Harb Perspect Biol. 2013;5.

29. Weyrich LS, Duchene S, Soubrier J, Arriola L, Llamas B, Breen J, et al. Neanderthal behaviour, diet, and disease inferred from ancient DNA in dental calculus. Nature. 2017;544: 357–361.

30. McGhee JJ, Rawson N, Bailey BA, Fernandez-Guerra A, Sisk-Hackworth L, Kelley ST. Meta-SourceTracker: application of Bayesian source tracking to shotgun metagenomics. PeerJ. 2020;8: e8783.

31. Cannon PF, Kirk PM. Fungal Families of the World. CABI; 2007.

32. Michelsen C, Winther Pedersen M, Fernandez-Guerra A, Zhao L, Petersen TC, Korneliussen TS. metaDMG – A Fast and Accurate Ancient DNA Damage Toolkit for Metagenomic Data. bioRxiv. 2022. p. 2022.12.06.519264. doi:10.1101/2022.12.06.519264

33. Maixner F, Krause-Kyora B, Turaev D, Herbig A, Hoopmann MR, Hallows JL, et al. The 5300-year-old Helicobacter pylori genome of the Iceman. Science. 2016;351: 162–165.

34. Rampelli S, Turroni S, Mallol C, Hernandez C, Galván B, Sistiaga A, et al. Components of a Neanderthal gut microbiome recovered from fecal sediments from El Salt. Commun Biol. 2021;4: 169.

35. Kjær KH, Winther Pedersen M, De Sanctis B, De Cahsan B, Korneliussen TS, Michelsen CS, et al. A 2-million-year-old ecosystem in Greenland uncovered by environmental DNA. Nature. 2022;612: 283–291.

36. Rognes T, Flouri T, Nichols B, Quince C, Mahé F. VSEARCH: a versatile open source tool for metagenomics. PeerJ. 2016;4: e2584.

37. Gutiérrez-García TA, Vázquez-Domínguez E, Arroyo-Cabrales J, Kuch M, Enk J, King C, et al. Ancient DNA and the tropics: a rodent’s tale. Biol Lett. 2014;10: 20140224–20140224.

38. Schroeder H, Ávila-Arcos MC, Malaspinas A-S, Poznik GD, Sandoval-Velasco M, Carpenter ML, et al. Genome-wide ancestry of 17th-century enslaved Africans from the Caribbean. Proc Natl Acad Sci U S A. 2015;112: 3669–3673.

39. Schroeder H, Sikora M, Gopalakrishnan S, Cassidy LM, Maisano Delser P, Sandoval Velasco M, et al. Origins and genetic legacies of the Caribbean Taino. Proc Natl Acad Sci U S A. 2018;115: 2341–2346.

40. Nägele K, Posth C, Iraeta Orbegozo M, Chinique de Armas Y, Hernández Godoy ST, González Herrera UM, et al. Genomic insights into the early peopling of the Caribbean. Science. 2020. doi:10.1126/science.aba8697

41. Nieves-Colón MA, Pestle WJ, Reynolds AW, Llamas B, de la Fuente C, Fowler K, et al. Ancient DNA Reconstructs the Genetic Legacies of Precontact Puerto Rico Communities. Mol Biol Evol. 2020;37: 611–626.

42. Fernandes DM, Sirak KA, Ringbauer H, Sedig J, Rohland N, Cheronet O, et al. A genetic history of the pre-contact Caribbean. Nature. 2021;590: 103–110.

43. Pochon Z, Bergfeldt N, Kirdök E, Vicente M, Naidoo T, van der Valk T, et al. aMeta: an accurate and memory-efficient ancient metagenomic profiling workflow. Genome Biol. 2023;24: 242.

44. Cribdon B, Ware R, Smith O, Gaffney V, Allaby RG. PIA: More accurate taxonomic assignment of metagenomic data demonstrated on sedaDNA from the North Sea. Front Ecol Evol. 2020;8. doi:10.3389/fevo.2020.00084

45. Astashyn A, Tvedte ES, Sweeney D, Sapojnikov V, Bouk N, Joukov V, et al. Rapid and sensitive detection of genome contamination at scale with FCS-GX. bioRxiv. 2023. doi:10.1101/2023.06.02.543519

46. Bálint B, Merényi Z, Hegedüs B, Grigoriev IV, Hou Z, Földi C, et al. ContScout: sensitive detection and removal of contamination from annotated genomes. Nat Commun. 2024;15: 936.

47. Uritskiy GV, DiRuggiero J, Taylor J. MetaWRAP-a flexible pipeline for genome-resolved metagenomic data analysis. Microbiome. 2018;6: 158.

48. Johnson M, Zaretskaya I, Raytselis Y, Merezhuk Y, McGinnis S, Madden TL. NCBI BLAST: a better web interface. Nucleic Acids Res. 2008;36: W5–9.

49. Warinner C, Herbig A, Mann A, Fellows Yates JA, Weiß CL, Burbano HA, et al. A Robust Framework for Microbial Archaeology. Annu Rev Genomics Hum Genet. 2017;18: 321–356.

50. Eisenhofer R, Cooper A, Weyrich LS. Reply to Santiago-Rodriguez et al.: proper authentication of ancient DNA is essential. FEMS microbiology ecology. 2017. doi:10.1093/femsec/fix042

51. Eisenhofer R, Weyrich LS. Proper Authentication of Ancient DNA Is Still Essential. Genes . 2018;9. doi:10.3390/genes9030122

52. Gilbert MTP, Hansen AJ, Willerslev E, Rudbeck L, Barnes I, Lynnerup N, et al. Characterization of Genetic Miscoding Lesions Caused by Postmortem Damage. Am J Hum Genet. 2003;72: 48–61.

53. Gilbert MTP, Binladen J, Miller W, Wiuf C, Willerslev E, Poinar H, et al. Recharacterization of ancient DNA miscoding lesions: insights in the era of sequencing-by-synthesis. Nucleic Acids Res. 2007;35: 1–10.

54. Briggs AW, Stenzel U, Johnson PLF, Green RE, Kelso J, Prüfer K, et al. Patterns of damage in genomic DNA sequences from a Neandertal. Proc Natl Acad Sci U S A. 2007;104: 14616–14621.

55. Jónsson H, Ginolhac A, Schubert M, Johnson PLF, Orlando L. mapDamage2.0: fast approximate Bayesian estimates of ancient DNA damage parameters. Bioinformatics. 2013;29: 1682–1684.

56. Neukamm J, Peltzer A, Nieselt K. DamageProfiler: fast damage pattern calculation for ancient DNA. Bioinformatics. 2021;37: 3652–3653.

57. Weiß CL, Dannemann M, Prüfer K, Burbano HA. Contesting the presence of wheat in the British Isles 8,000 years ago by assessing ancient DNA authenticity from low-coverage data. Elife. 2015;4. doi:10.7554/eLife.10005

